# LDscore: a scalable, Python 3-powered web platform for LD score regression analysis

**DOI:** 10.64898/2025.12.19.695639

**Authors:** Charles E. Breeze, Xiaozheng Yao, Brian Park, Madhulika Kanigicherla, Qing Lan, Nathaniel Rothman, Andrew E. Teschendorff, Nora Franceschini, Sonja I. Berndt, Mitchell J. Machiela

## Abstract

Linkage disequilibrium score regression (LDSC) is an important analytical tool for quantifying heritability and estimating genetic correlations between complex traits. However, the LDSC original implementation relies on an outdated Python 2 framework and deploying the standard command-line tools requires significant setup, data access, and computational expertise, creating a barrier for many researchers. To overcome these limitations, we developed LDscore, a significant technical and accessibility upgraded version of LDSC that allows for rapid analysis of GWAS data. The core advancement is the recoding of the LDSC framework in Python 3, enabling computational optimization and ensuring long-term sustainability. Built on top of this improved foundation, LDscore is implemented as a free, publicly available web application integrated within the popular NCI LDlink framework. LDscore can accelerate scientific research by providing an intuitive graphical interface for heritability estimation, genetic correlation, and LD score calculation, including access to an expanded range of reference populations for online analysis. Notably, our results show that selecting the most appropriate reference population LD panel, even at the subcontinental ancestry group level, is essential for minimizing population stratification bias in heritability estimation. By leveraging cloud computing for superior scalability and eliminating the need for local installation, LDscore adheres to FAIR principles, improving access, traceability, and reproducibility across an expanded set of reference populations, and effectively widens access to researchers worldwide providing support for in-depth genetic analyses.

**Brief summary:** Linkage disequilibrium score regression (LDSC), a widely-used method for quantifying heritability and genetic correlation, is limited by an outdated Python 2 framework and complex command-line deployment. We developed LDscore, a significant technical upgrade built on Python 3 for sustainability and computational optimization. LDscore is a free, cloud-based web application integrated into NCI LDlink. LDscore eliminates installation barriers, offering an intuitive interface for computing heritability estimates, LD scores, and genetic correlation. Crucially, LDscore expands the range of reference populations available in LDSC, which can reduce population-stratification-based bias. Leveraging cloud computing, LDscore accelerates and widens global researcher access to LDSC-based genetic computation.

**Availability:** LDscore is freely available within LDlink at https://ldlink.nih.gov/ldscore. Source code for the updated LDSC Python3 framework is available at https://github.com/CBIIT/ldsc under the GNU General Public License v3.0 and the webtool code is at https://github.com/CBIIT/nci-webtools-dceg-linkage (webtool code) under the MIT license.

## Background

Genome-wide association studies (GWAS) have successfully identified thousands of loci associated with complex human traits [1] [2] [3]. Linkage disequilibrium score regression (LDSC) provides a robust framework for interpreting these results by helping to distinguish polygenicity from confounding factors like population stratification, estimating SNP-heritability, evaluating genetic correlations with other diseases, traits, and annotations, and providing support for further analyses using GWAS summary statistics [4] [5].

Despite its valuable contribution to the interpretation of GWAS, the utility of the published LDSC toolkit has become increasingly limited by its dependency on a deprecated Python 2 environment. This Python 2 requirement poses significant security and maintenance challenges. It often necessitates complex system setups to limit vulnerability, may not be permissible at some institutions, and prevents many researchers from readily utilizing the method. In addition, precomputed population linkage disequilibrium (LD) score reference panels in the original toolkit are limited to Europeans and East Asians from the 1000 Genomes Project[6], making it difficult to utilize the method for other populations with different LD structures.

Our tool, LDscore, addresses these limitations by first providing a massive technical overhaul of the method, migrating the core LDSC framework to Python 3. This modernization ensures greater long-term code sustainability, reduces security vulnerability concerns, and enables subsequent improvements in speed and computational optimization. We have also created a more extensive set of precomputed LD references from a broad range of populations from both the 1000 Genomes Project [6] and Genome Aggregation Database (gnomAD) [7]. Built upon this robust new framework, LDscore is delivered as a user-friendly, cloud-based web application. This approach eliminates the historical requirement for complex environment setup, Python version access, and specialized computational resources. This commitment to accessibility effectively broadens its use for researchers globally, providing a powerful, stable, and ready-to-use platform for high-quality genetic analysis.

## Results and Discussion

The LDscore web application is integrated into LDlink, which is a widely used, web-based suite of genetics programs designed to aid researchers in visualizing and exploring LD structure in different populations [8]. The LDscore web application facilitates three core analyses (**Figure 1A**): 1) SNP-heritability estimation (h^2^_SNP_) and associated statistics, including the genomic inflation factor (λ_GC_), LDSC intercept (I_LDSC_) and attenuation ratio statistic (R), which are important for assessing population stratification [4,9]; 2) genetic correlation, which allows for the estimation of the genetic overlap between two user-uploaded GWAS summary statistics files; and 3) LD score calculation, which provides rapid computation of LD scores of all SNPs, including summaries by minor allele frequency for given dataset in PLINK [10] format (e.g.,*.bed, *.bim, and *.fam files). To ensure reproducibility, we compared LD score regression estimates for the heritability between the original Python 2 implementation and LDscore Python 3 implementations for a GWAS meta-analysis on rheumatoid arthritis [11] [12]. Results show perfect agreement between these two implementations (**Figure 1B**), as well as consistency with estimates from the literature [13].

**Figure 1:**
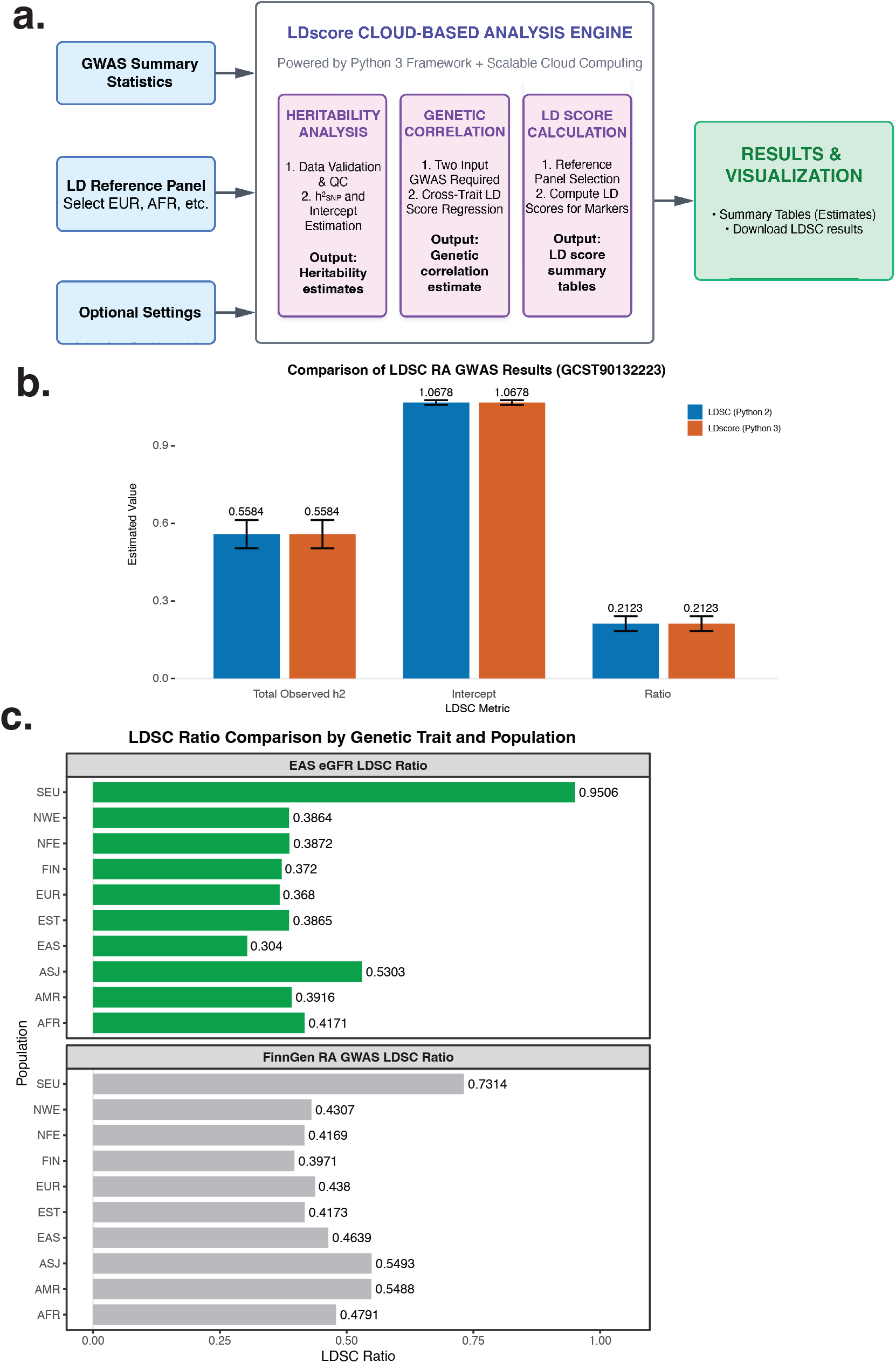
LDscore Web Application: An Integrated and Updated Workflow for LD Score Regression Analysis. **[a]** LDscore Key Feature: All analyses are performed via a modern, scalable Python 3 engine on cloud infrastructure, eliminating the need for complex local installation or command-line expertise. **[b]** Comparison of rheumatoid arthritis (RA) GWAS results between LDSC Python2 and LDscore Python3 shows reproducibility across heritability (h2), intercept and ratio LDSC metrics, as well as standard errors (error bars). **[c]** Comparison of LDSC Ratios across ten global populations. Bar plots represent results for EAS estimated glomerular filtration rate (eGFR) (top panel), and FinnGen Rheumatoid Arthritis GWAS (bottom panel). The X-axis indicates the LDSC Ratio, and the Y-axis lists the population group.

One of the core analysis options in LDSC is selection of the population-specific pre-indexed LD reference panel. We sought to empirically assess the impact of reference panel choice on LDSC estimation bias. To do this, we analyzed data from a FinnGen GWAS on rheumatoid arthritis (RA) and a BioBank Japan GWAS on estimated glomerular filtration rate (eGFR). For both traits, analysis using the corresponding population-specific reference (FIN and EAS, respectively) yielded the lowest LDSC attenuation ratio, which indicates minimized population stratification bias (**Figure 1C**). This observation underscores the requirement for population-matched LD references to accurately mitigate bias in LDSC-based analyses. Consequently, we expanded the pre-indexed LD reference panels in LDscore to cover 10 distinct reference populations. These additional pre-indexed LD reference panels were derived from large, high-quality, publicly available genomic datasets, including the 1000 Genomes Project and the Genome Aggregation Database (gnomAD) (**Table 1, Table S1**) [6] [7]. Rather than requiring users to find and download a LD reference panel or generate their own LD reference data, a computationally intensive and error-prone process, LDscore provides immediately accessible LD reference panels. The precomputed LD reference panels provided in LDscore are based on high-density marker information with substantial quality control, allowing for accurate LD score analyses. These panels provide increased representation and allow better LD matching with a broader range of GWAS summary statistics and populations, ultimately resulting in improved estimates of LDSC parameters.

**Table 1.** List of populations reference LD panels included with LDSC and LDscore.

| **Population LD reference panel** | **Original LDSC (Python2)** | **LDscore (Python3)** |
| --- | --- | --- |
| a. (EUR) European | Yes | Yes |
| b. (EAS) East Asian | Yes | Yes |
| c. (AFR) African American | No | Yes |
| d. (ASJ) Ashkenazi Jewish | No | Yes |
| e. (EST) Estonian | No | Yes |
| f. (FIN) Finnish European | No | Yes |
| g. (AMR) Latino/Admixed American | No | Yes |
| h. (NFE) Non-Finnish European | No | Yes |
| i. (NWE) North-Western European | No | Yes |
| j. (SEU) Southern European | No | Yes |

To utilize heritability and genetic correlation in LDscore, users need to provide a set of GWAS summary statistics; LD score calculation requires individual level genotype data. Once the dataset is uploaded, a data checking step is performed to ensure the uploaded summary statistics adhere to necessary formatting standards for the analysis, minimizing common user errors. Example datasets are also included on the webtool to allow the user to see how the program works before uploading their own dataset (a walk-through example is also available in the **Appendix**). Upon analysis completion, LDscore generates interactive summary tables, including LD score mean, standard deviation, min, Q1, Q3, and max, and heritability and genetic correlation statistics, all of which are available for download in tabular format.

Our cloud-based webtool provides an easy-to-use framework with minimal data formatting. Users benefit from a tangible increase in speed of computational setup, with typical setup taking a few minutes (GUI training). Previous implementations of LDSC often involved extensive setup, including coding and scripts to perform the analysis and overcome computational bottlenecks. Our cloud implementation allows researchers to immediately focus on the scientific analysis and interpretation of their results. The pre-processed LD reference panels remove a substantial technical barrier, allowing researchers to rapidly apply LD-score regression to their summary statistics with appropriately matched LD information, thus streamlining large-scale genomic analyses. Furthermore, by providing a broad range of LD reference panels and computation, LDscore significantly improves access, traceability, and reproducibility, demonstrating adherence to FAIR principles and promoting better, more accessible science [14].

## Conclusions

LDscore upgrades the Linkage Disequilibrium Score Regression (LDSC) analytical framework by recoding LDSC in Python 3. This technical advance ensures long-term code sustainability, security, and computational optimization. Delivered as a scalable, cloud-based web application within NCI LDlink, LDscore effectively eliminates the primary barriers to utilizing LDSC -namely, the requirement for complex local installation and specialized computational expertise. The platform provides a user-friendly interface for SNP-heritability estimation, genetic correlation analysis, and LD score calculation. Critically, LDscore includes an expanded set of pre-indexed LD reference panels, thus increasing the accuracy and applicability of LDSC to diverse populations. By providing immediate access to a validated and reproducible analytical engine, LDscore adheres to FAIR principles, significantly promoting the accessibility and reproducibility of genetic analyses worldwide.

## Availability

LDscore is an open-source project and is accessible without registration at https://ldlink.nih.gov/ldscore. A detailed user guide and a frequently asked questions (FAQ) section are provided at https://ldlink.nih.gov/help#LDscore to assist new users. The underlying source code, which includes the front-end interface and the deployment scripts for the LDSC container environment, is maintained on GitHub at https://github.com/cbiit/ldsc (updated LDSC in Python 3) under the GNU General Public License v3.0 and https://github.com/CBIIT/nci-webtools-dceg-linkage (webtool code) under the MIT license.

## Methods

### 1. Aim, Design, and Setting

The aim was to upgrade the Linkage Disequilibrium Score Regression (LDSC) toolkit by recoding the core framework from Python 2 to Python 3, thus enhancing computational sustainability and performance. In addition, we also developed LDscore, a scalable web application that provides easy access to LDSC analysis, including access to LD references across diverse populations within the secure NCI cloud computing environment. The tool is publicly available within the LDlink web suite at https://ldlink.nih.gov/ldscore.

### 2. Materials

#### Input Materials

The platform is designed to process user-provided genome-wide association study (GWAS) summary statistics. Heritability and genetic correlation calculation require summary statistics from GWAS. LD score calculation requires individual-level genotype data in the standard PLINK format (*.bed, *.bim, and *.fam files).

#### Reference Materials

Analysis relies on an expanded set of pre-computed linkage disequilibrium (LD) reference panels across diverse populations derived from the 1000 Genomes Project and the genome aggregation database (gnomAD). These panels provide expanded population coverage for appropriate LD matching.

### 3. Processes and Interventions

#### System Architecture

The LDscore application is delivered via the popular LDlink interface, employing a hybrid web stack (HTML, CSS, JavaScript frontend) with a Python Flask backend. The core analytical engine, the modernized Python 3 LDSC framework, is containerized and executed within the robust NCI cloud computing infrastructure. This cloud-based approach eliminates the requirement for local software installation or dependency management for the end-user.

#### Analytical Workflow

The workflow is initiated by user upload of one or more GWAS summary statistics files:

Validation and Pre-processing: Uploaded data undergoes automated validation to check for correct header conventions and alignment with the required LDSC format.

Execution: Jobs are queued and executed using the Python 3 LDSC engine on the cloud. The infrastructure manages parallel jobs, ensuring high throughput and employing appropriate data handling protocols.

Output: Computed results, including summary tables, output logs, and statistical estimates, are rendered directly in the browser and are available for download in tabular format.

### 4. Statistical Analysis and Software Requirements

#### Statistical Analysis

The platform performs Linkage Disequilibrium Score Regression (LDSC). Key statistical outputs include:

SNP-Heritability Estimation (h^2^_SNP_).

Genetic Correlation between uploaded summary statistics.

Confounding Assessment via the genomic inflation factor (lambda_GC), LDSC intercept, and attenuation ratio statistic.

LD Score Computation for individual SNPs.

#### Validation and Software Requirements

Method Validation: Equivalence was established by comparing LDSC parameter estimates (including h^2^_SNP_ and LDSC intercept) for a publicly available rheumatoid arthritis GWAS dataset between the new Python 3 LDscore implementation and the original Python 2 implementation. The comparison demonstrated perfect agreement (**Figure 1B**).

Software Tool Requirements: LDscore is delivered as an easy-to-use web application; no local software dependencies are required for the end-user. The source code for the LDSC framework is available under the GNU General Public License v3.0, and the web tool code is under the MIT license.

## Acknowledgements

This research was supported by the Intramural Research Program of the National Institutes of Health (NIH). The contributions of the NIH authors were made as part of their official duties as NIH federal employees, are in compliance with agency policy requirements, and are considered Works of the United States Government. However, the findings and conclusions presented in this paper are those of the authors and do not necessarily reflect the views of the NIH or the U.S. Department of Health and Human Services.

## Contributions

Advised or performed statistical analyses (CEB, XY, BP, MK, QL, NR, AT, NF, SIB, MJM) Web tool design (CEB, XY, BP, MK, SIB, MJM) CEB and MJM wrote the paper with contributions from all other authors. All authors contributed to the final draft.

## Data availability

Rheumatoid arthritis GWAS meta-analysis data are available from https://www.ebi.ac.uk/gwas/studies/GCST90132223

FinnGen rheumatoid arthritis GWAS data are available from https://opengwas.io/datasets/finn-b-M13_RHEUMA#

BioBank Japan estimated glomerular filtration rate GWAS data are available from https://opengwas.io/datasets/ieu-a-1284

European LD reference data are available at https://zenodo.org/records/8182036

LD reference data across other populations are available at http://gnomad-sg.org/downloads#v2-linkage-disequilibrium

---

**Table S1.** Additional information for populations covered in LDscore webtool reference LD panels (more information for 1000 genomes is available at https://grch37.ensembl.org/info/genome/variation/species/populations.html and for gnomAD at https://gnomad.broadinstitute.org/news/2018-10-gnomad-v2-1/).

| **Population** | **Description** | **Genomes** | **Source** |
| --- | --- | --- | --- |
| AFR | African/African American | 4,359 | gnomAD |
| AMR | Latino/Admixed American | 424 | gnomAD |
| ASJ | Ashkenazi Jewish | 145 | gnomAD |
| FIN | Finnish | 1,738 | gnomAD |
| NFE | Non-Finnish European | 7,718 | gnomAD |
| EST | Estonian | 2,297 | gnomAD |
| NWE | North-Western European | 4,299 | gnomAD |
| SEU | Southern European | 53 | gnomAD |
| EAS | East Asian | 780 | gnomAD |
| EUR | European | 503 | 1000 genomes |

## Appendix

### Walk-through Example of LDscore (Tutorial)

The LDscore web application is designed to guide the user through the process of LD score regression analysis with minimal setup. We provide a step-by-step tutorial using publicly available summary statistics from the Biobank Japan (BBJ) GWAS for high-density lipoprotein (HDL) cholesterol levels (file BBJ.HDL-C.autosome.txt.gz, available from http://jenger.riken.jp/47analysisresult_qtl_download/). A subset of these data for chromosome 22 are also available via the “Use example data” toggle at https://ldlink.nih.gov/ldscore (file BBJ_HDLC22.txt).

LDscore consists of three core modules: 1) Heritability Estimation, 2) Genetic Correlation, and 3) LD Score Calculation. These modules are accessed via distinct tabs within the unified LDlink interface.

#### A.1. Heritability Analysis (Figure S1)

The first module, Heritability Analysis, allows users to estimate the SNP-heritability (h^2^_SNP_) for a given trait or disease as well as other metrics, including the genomic inflation factor, LDSC intercept and attenuation ratio statistic, to help assess the relative contributions of polygenicity and population stratification to the GWAS summary statistics.

1. Select Tab: Navigate to the “Heritability Analysis” tab and select.
2. Data Upload: Upload a GWAS summary statistics file (e.g., BBJ.HDL-C.autosome.txt.gz). Details regarding the required format for GWAS summary statistics can be found by clicking on the sample format. LDscore automatically initiates a data checking step to ensure the file adheres to the required format (e.g., correct header columns, such as SNP, A1, A2, N, Z). An error message will be returned if GWAS summary statistics are formatted incorrectly for analysis.
3. Reference Panel Selection: Select the reference panel that most closely resembles the population used for the GWAS summary statistics from the dropdown menu. This is a crucial step to ensure that the LD scores are calculated using the appropriate population-specific linkage disequilibrium scores, minimizing bias and error. The example provided is from a Japanese cohort, so the user should select the (EAS) East Asian population from the dropdown menu for the example.
4. Perform Analysis: Click the “Calculate” button. The job will be queued on the NCI cloud infrastructure.
5. Display of Results (Figure S1): Upon completion, the page displays the results summary. The user can view the:
  - Total observed scale h^2^ (h^2^_SNP_): LDSC-estimated SNP heritability estimate and standard error for the disease of trait.
  - Lambda GC (λ_GC_): Genomic inflation factor for GWAS summary statistics
  - Mean chi2: mean chi-square value from GWAS summary statistics
  - LDSC Intercept (I_LDSC_): Intercept value from LDSC analysis.
  - Ratio (R): Attenuation ratio statistic, which may provide a better quantification of population stratification.

#### A.2. Genetic Correlation (Figure S2)

This module enables the estimation of the genetic correlation between two distinct traits or diseases using LDSC. In this example, we correlate HDL cholesterol with another lipid trait, low-density lipoprotein (LDL) cholesterol (available from http://jenger.riken.jp/61analysisresult_qtl_download/).

1. Select Tab: Navigate to the “Genetic Correlation” tab and select.
2. Data Upload: For this module, two separate GWAS summary statistics files must be uploaded, one for each trait or disease (e.g., files BBJ_HDLC22.txt and BBJ_LDLC22.txt in example set). Details regarding the required format for GWAS summary statistics can be found by clicking on the sample format. LDscore automatically initiates a data checking step to ensure the file adheres to the required format (e.g., correct header columns, such as SNP, A1, A2, N, Z). An error message will be returned if GWAS summary statistics are formatted incorrectly for analysis.
3. Reference Panel Selection: Select the reference panel that most closely resembles the populations used for both GWAS summary statistics from the dropdown menu. Again, for this example, the (EAS) East Asian population should be select, because the GWAS summary statistics come from the BioBank Japan cohort.
4. Perform Analysis: Click the “Calculate” button. The platform will execute the cross-trait LD Score Regression.
5. Display Results (Figure S2): The resulting output gives the estimated genetic correlation between the two traits or disease and its associated p-value. A significant positive result suggests shared genetic architecture. The module also reports the h^2^_SNP_ estimate for each individual trait as a quality check.

#### A.3. LD Score Calculation (Figure S3)

This module allows users to generate LD scores for custom genomic annotations or subsets of markers, providing maximum flexibility for specialized analyses.

1. Select Tab: Navigate to the “LD Score Calculation” tab and select.
2. Data Upload: For this analysis, PLINK-formatted files (e.g. *.bed, *.bim, *.fam) consisting of primary genetic data are required, allowing for the rapid computation of LD scores and associated summaries. If the data are formatted incorrectly for analysis, an error message will appear.
3. Perform Analysis: Click the “Calculate” button. The application calculates the LD scores for the dataset.
4. Displayed Results (Figure S3): The primary output is a summary table detailing the computed LD scores, along with key descriptive statistics of the LD scores generated (mean, standard deviation, min, Q1, Q3, and max). This output can then be used in custom analyses or other downstream applications. All computed LD scores are provided as a downloadable file, ensuring the work adheres to FAIR standards. All genotype data used for this analysis is deleted from the server after analysis.

**Figure S1.**
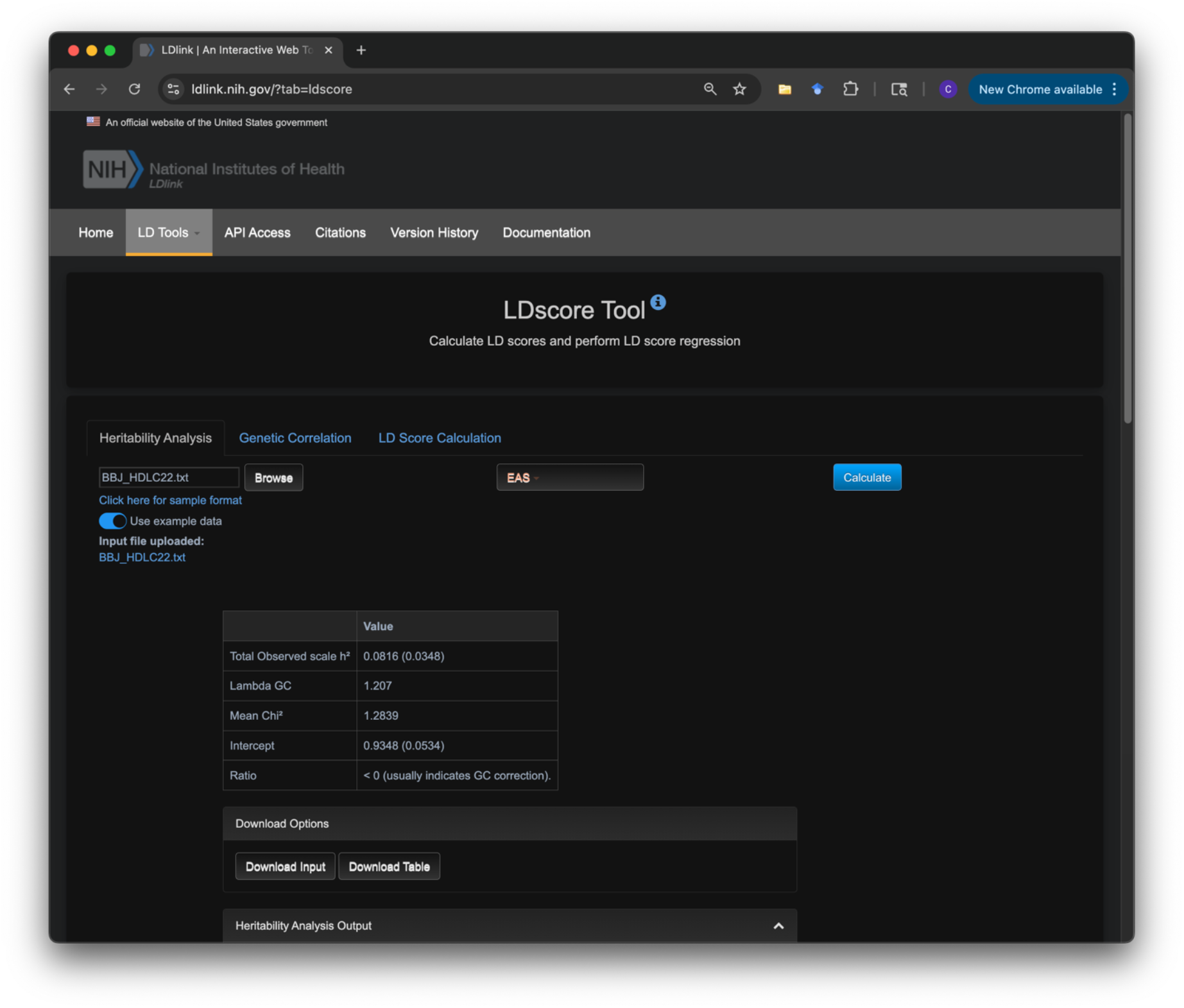
LDscore webtool, with an example Biobank Japan HDL heritability analysis (using the reference East Asian -EAS-panel). Note the tabs for “Heritability Analysis”, “Genetic Correlation”, and “LD Score Calculation”.

**Figure S2.**
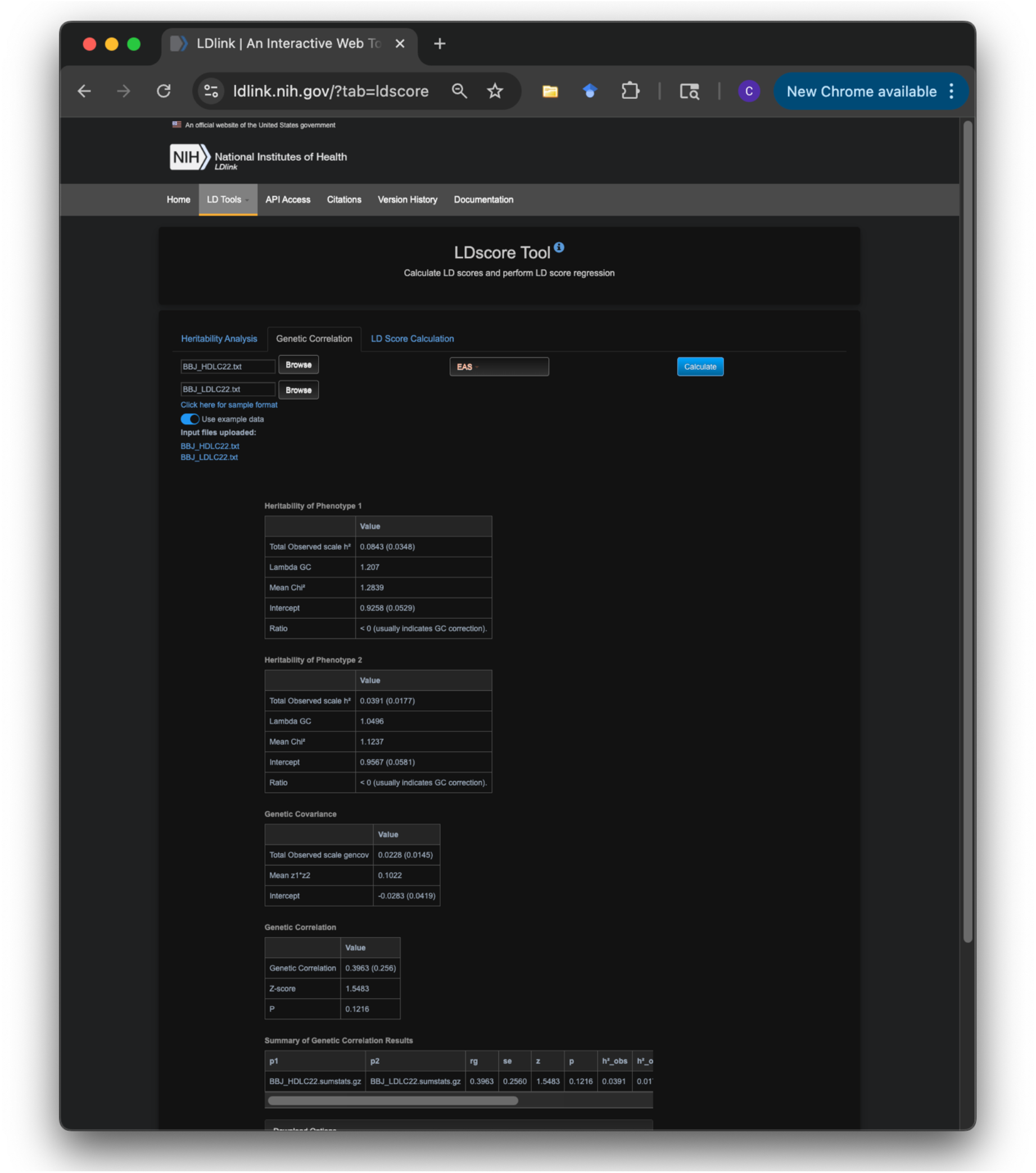
LDscore webtool, with an example Biobank Japan HDL genetic correlation analysis (using the reference East Asian -EAS-panel). Note the tabs for “Heritability Analysis”, “Genetic Correlation”, and “LD Score Calculation”.

**Figure S3.**
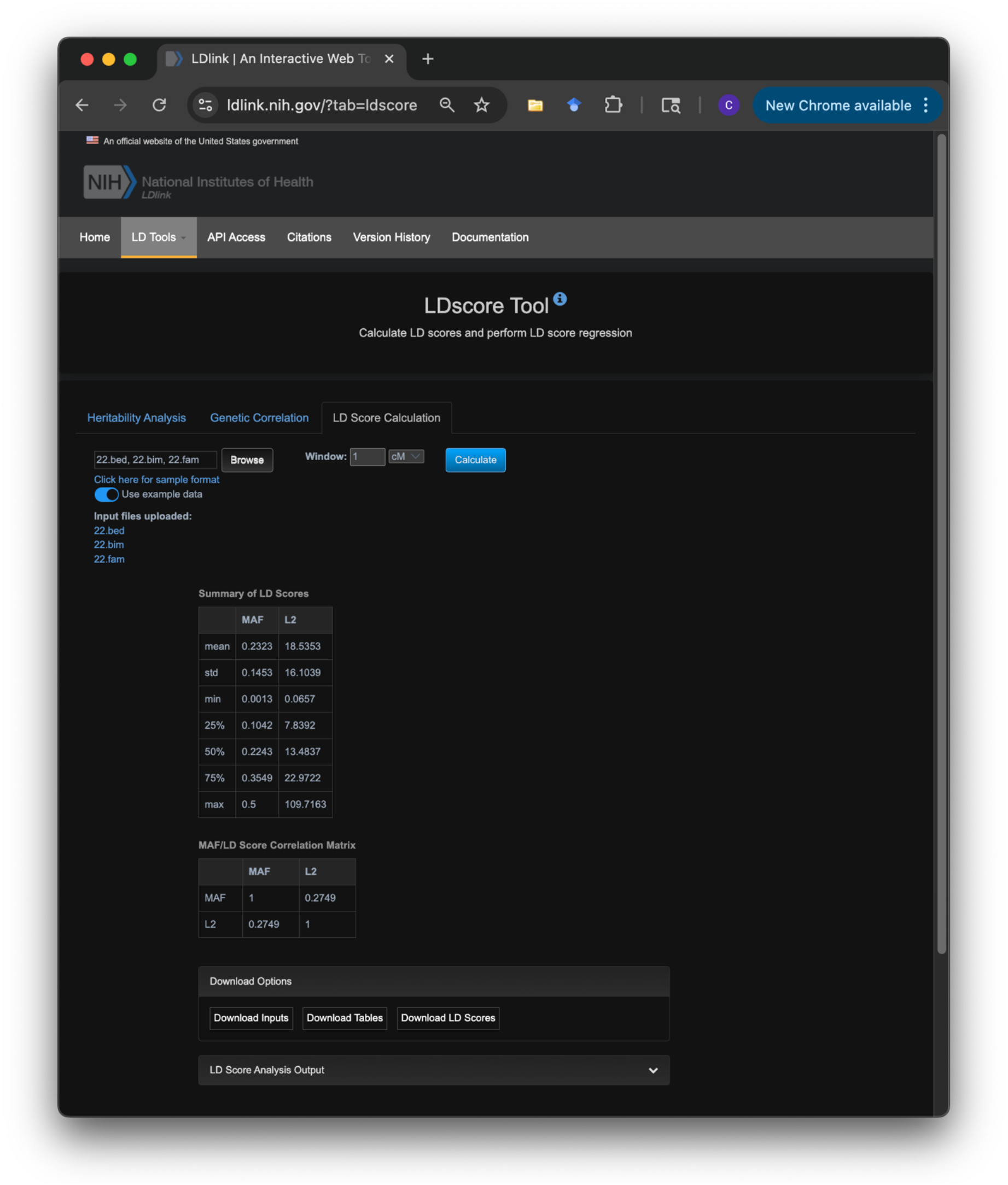
LDscore webtool, with an example Biobank Japan HDL LD score calculation. Note the tabs for “Heritability Analysis”, “Genetic Correlation”, and “LD Score Calculation”.

